# Molecular dissection of neurodevelopmental disorder-causing mutations in CYFIP2

**DOI:** 10.1101/2020.02.11.943332

**Authors:** Matthias Schaks, Klemens Rottner

**Affiliations:** Division of Molecular Cell Biology, Zoological Institute, Technische Universität Braunschweig, Spielmannstrasse 7, 38106 Braunschweig, Germany; Department of Cell Biology, Helmholtz Centre for Infection Research, Inhoffenstrasse 7, 38124 Braunschweig, Germany; Braunschweig Integrated Centre of Systems Biology (BRICS), Rebenring 56, 38106 Braunschweig, Germany

**Keywords:** WAVE regulatory complex, Arp2/3, protrusion, lamellipodium, CRISPR, Cas9

## Abstract

Actin remodelling is frequently regulated by antagonistic activities driving protrusion and contraction downstream of Rac and Rho small GTPases, respectively. WAVE regulatory complex (WRC), which primarily operates downstream of Rac, plays pivotal roles in neuronal morphogenesis. Recently, two independent studies described de novo mutations in the CYFIP2 subunit of WRC, which caused intellectual disability (ID) in humans. Although mutations had been proposed to effect WRC activation, no experimental evidence for this was provided. Here, we made use of CRISPR/Cas9-engineered B16-F1 cell lines that were reconstituted with ID-causing CYFIP variants in the context of compromised WRC activation with or without reduced Rac activities, which established that the majority of CYFIP2 mutations (5 out of 8) indeed cause constitutive WRC activation. Strikingly, activating mutations are positioned in a conserved WAVE- binding region mediating WRC transinhibition. As opposed to such gain-of-function mutations, a truncating mutant represented a loss-of-function variant, because it failed to interact with WRC components, and two mutants displayed no or at best a moderate increase of WRC activation. Collectively, our data show that CYFIP2 mutations frequently but not always coincide with WRC activation and suggest that normal brain development requires a delicate and precisely tuned balance of neuronal WRC activity.

## 1. Introduction

Branched actin filament networks, created by Actin-related protein 2/3 (Arp2/3) complex, play crucial roles in cell motility, neuronal path finding and morphogenetic events such as dendrite branching [1–5]. The heteropentameric WAVE regulatory complex (WRC) is the major Arp2/3 complex activator in protrusive structures such as lamellipodia, growth cones or dendrite branchlets, each mediating distinct, aforementioned processes [3,4,6–8]. WRC itself is controlled by a variety of different signalling inputs, but binding to the small GTPase Rac appears to be the most crucial one [2,7,9,10]. While initially proposed to disassemble into two subcomplexes [11], more recent literature strongly suggests that Rac binding to WRC releases the trans-inhibitory interaction of the Arp2/3 complex-activating WCA domain of WAVE with CYFIP1/2 (see below), but without WRC subunit dissociation [7,9,10]. WRC is composed of five different components: WAVE2 (or the paralogs WAVE1/WAVE3), CYFIP1 – also called Sra-1 (or the paralog CYFIP2), Nap1 (or the hematopoietic paralog Hem1), Abi1 (or Abi2/3) and HSPC300 [11–14]. The CYFIP subunit is the regulatory subunit within WRC. CYFIP is on the one hand sequestering WAVE’s WCA domain and on the other hand receiving signalling input from Rac by binding through two distinct GTPase binding sites, called A and D site, for adjacent and distant to the WCA binding site, respectively. Rac binding is commonly agreed to release the WCA domain from transinhibition [10,15]. Due to the extremely high sequence conservation in respective regions, it is assumed that CYFIP1 and CYFIP2 function entirely redundantly with respect to WRC activation.

Evidence for a relevance of WRC and thus Arp2/3-dependent actin remodelling in different tissues is continuously growing [3,4,6,8,16,17]. This view is complemented by the discovery of neurodevelopmental disorders, which coincide with genetic aberrations in the *CYFIP1* locus. Deletions involving 15q11–q13 also harbouring the *CYFIP1* locus, are relatively common. Many of these rearrangements are associated with abnormal phenotypes including seizure, developmental delay and autism, but deletions affecting *CYFIP1* typically cause a worse phenotype, compared to deletions in these regions not involving CYFIP1 [18]. Much more direct, however, are recent studies showing *de novo* mutations in the Rac / WAVE regulatory complex (WRC) pathway to be causative for neurodevelopmental disorders and intellectual disabilities. Two studies found mutations in the *NCKAP1* gene, encoding for Nap1, with unknown functions [19,20]. While loss-of-function mutations have been described for the *WASF1* gene [21], encoding the protein WAVE1, another recent study found mutations in the *RAC1* gene and suggested these mutations to either generate dominant negative or constitutively active alleles [22]. Other studies found mutations in *CYFIP2*, which were accused to cause gain-of-function with respect to WRC activation [23,24], but experimental evidence for this assumption was hitherto missing.

As a model system to systematically analyse the molecular consequences of CYFIP2 mutations normally occurring in neurodevelopmental disorders, we focused on B16-F1 mouse melanoma cells, in which the *CYFIP1* and *CYFIP2* genes were disrupted using CRISPR/Cas9 [7]. CYFIP1/2 removal causes complete failure to form Rac-dependent lamellipodia, which can be readily restored by ectopic expression of CYFIP1. These Arp2/3 complex-rich, lamellipodial actin networks constitute the best-characterised, WRC-dependent structures, but they also display high similarity to growth cones [6]. We propose that results obtained with this cell-based, morphological assay can be directly translated into functions of WRC in similar structures, such as neuronal growth cone or dendrite branchlet common to the nervous system.

## 2. Materials and Methods

### Cell Culture

B16-F1 cell line was purchased from ATCC (CRL-6323, sex:male). B16-F1 derived CYFIP1/2 KO cells (clone #3) were as described. B16-F1 cells and derivatives were cultured in DMEM (4.5 g/l glucose; Invitrogen), supplemented with 10% FCS (Gibco), 2 mM glutamine (Thermo Fisher Scientific) and penicillin (50 Units/ml)/streptomycin (50 µg/ml) (Thermo Fisher Scientific). B16-F1 cells were routinely transfected in 35 mm dishes (Sarstedt), using 0.5 µg DNA in total and 1 µl JetPrime for controls, and 1 µg DNA in total and 2 µl JetPrime for B16-F1-derived knockout cells. After overnight transfection, cells were plated onto acid-washed, laminin-coated (25 µg/ml) coverslips and allowed to adhere for at least 5 hours prior to analysis.

### DNA constructs

pEGFP-C2 vector was purchased from Clontech Inc. (Mountain View, CA, USA). pEGFP-C2-Sra-1 (CYFIP1), and derived mutant contructs (i.e. A site [C179R/R190D] and WCA* [L697D/Y704D/L841A/F844A/W845A]) were described previously [7] and correspond to the splice variant *CYFIP1a*, sequence AJ567911. CYFIP2-related mutations were introduced in the respective positions in CYFIP1 by site-directed mutagenesis, based on sequence alignment (Figure S1). The identity of DNA constructs was verified by sequencing.

### CRISPR/Cas9-mediated genome editing

B16-F1 cells lacking functional *CYFIP1* and *CYFIP2* genes, as well as reduced expression of Rac GTPases were generated by treating CYFIP1/2 KO cells (clone #3) with pSpCas9(BB)-2A-Puro (PX459) vectors targeting Rac1, Rac2 and Rac3 genes, as described (REF). Specifically, cells were co-transfected with plasmids targeting ATGCAGGCCATCAAGTGTG (Rac1/2) and ATGCAGGCCATCAAGTGCG (Rac3) genomic regions as described [7]. After puromycin selection of transfected cells (3 days), cells were extensively diluted and a few days later, macroscopically visible colonies picked, to obtain single cell-derived clones. Derived cell clones already lacking CYFIP1/2 were screened for low expression of Rac GTPases by Western Blotting.

### Western blotting

Preparation of whole cell lysates was performed essentially as described [7]. Western blotting was carried out using standard techniques. Primary antibodies used were CYFIP1/2 (Sra-1/PIR121 [14], Rac1/3 (23A8, Merck), Nap1 [14], WAVE [7], Abi1 [7] and GAPDH (6C5, Calbiochem). HRP-conjugated secondary antibodies were purchased from Invitrogen. Chemiluminescence signals were obtained upon incubation with ECL™ Prime Western Blotting Detection Reagent (GE Healthcare), and were recorded with ECL Chemocam imager (Intas, Goettingen, Germany).

### Immunoprecipitation

For EGFP-immunoprecipitation experiments, B16-F1 cells expressing EGFP or EGFP-tagged variants of CYFIP1 were lysed with lysis buffer (1% Triton X-100, 140 mM KCl, 50 mM Tris/HCl pH 7.4/50 mM NaF, 10 mM Na_4_P_2_O_7_, 2 mM MgCl_2_ and Complete Mini, EDTA-free protease inhibitor [Roche]). Lysates were cleared and incubated with GFP-Trap agarose beads for 60 min. Subsequently, beads were washed three times with lysis buffer lacking protease inhibitor, mixed with Laemmli buffer, boiled for 5 min and subjected to Western Blotting.

### Fluorescence microscopy, phalloidin stainings and quantification

B16-F1-derived cell lines were seeded onto laminin-coated (25 µg/ml), 15 mm-diameter glass coverslips and allowed to adhere for at least 5 hours. Cells were fixed with pre-warmed, 4% paraformaldehyde (PFA) in PBS for 20 min, and permeabilized with 0.05% Triton-X100 in PBS for 30 sec. PFA-fixed cell samples following transfections with plasmids mediating expression of EGFP-tagged proteins were counterstained with ATTO-594-conjugated phalloidin.

### Time-lapse microscopy

Live cell imaging was done with CYFIP1/2 KO cells (clone #3) transfected with respective EGFP-tagged CYFIP1 variants and migrating on laminin-coated glass (25 µg/ml). Cells were observed in µ-Slide 4 well (Ibidi), and maintained in microscopy medium (F12 HAM HEPES-buffered medium, Sigma), including 10% FCS, 2 mM L-glutamine and penicillin (50 Units/ml)/streptomycin (50 µg/ml) (Thermo Fisher Scientific). Conventional video microscopy was performed on an inverted microscope (Axiovert 100TV, Zeiss) equipped with an HXP 120 lamp for epifluorescence illumination, a halogen lamp for phase-contrast imaging, a Coolsnap-HQ2 camera (Photometrics) and electronic shutters driven by MetaMorph software (Molecular Devices). Live cell images were obtained using a 100 x/1.4 NA Plan apochromatic oil objective at a frame rate of 12/min. Kymographs were generated using MetaMorph software by drawing lines from inside the cell and across the cell edge, and the protrusion velocity determined by measuring the advancement of cell edges over time.

### Calculation of surface conservation

Amino acid conservation scores of CYFIP1 within WRC (PDB ID: 3P8C) were calculated using the ConSurf webserver [25]. A total of 186 unique protein sequences ranging from 30 to 95 % identity were used to calculate conservation (scored from 1 to 9) and surface conservation colour-coded as indicated in the figure.

### Statistical analysis

To assess statistical significance, two-sided, two-sample t-test was applied when data passed normality and equal variance tests, otherwise nonparametric Mann-Whitney-Rank-Sum test was performed. Statistical analyses were performed using Sigma plot 12.0 (Systat Software). Observed differences between groups were considered to be statistically significant if the error probability (p-value) of this assumption was below 5 % (p < 0.05).

## 3. Results

Our previous results demonstrated that B16-F1 cells disrupted for both CYFIP1 and −2 expression (Sra-1/PIR121-KO clone #3) lack lamellipodia entirely, but can be restored to form lamellipodia upon expression of CYFIP1 wildtype [7]. It was also established previously that mutational inactivation of one of the two Rac-binding site in CYFIP1, the so called A site, abrogated WRC activation, since A site-mutated CYFIP1 was almost entirely abolished in its capability to restore lamellipodia formation ([7] and Figure 1A). This defect, however, was completely eliminated upon additional mutation of CYFIP1 residues essential for binding to the WCA-domain of WAVE proteins, hence a modification rendering WRC constitutively active and termed WCA* [7,10]. These data unequivocally established that the major role of the A site interacting with Rac is in activation of WRC to drive lamellipodial edge localization and Arp2/3 complex activation [7]. To ask whether previously described CYFIP2 mutations found in human patients might indeed also render WRC constitutively active, we transferred these mutations to A site-mutated, murine CYFIP1. Note that murine CYFIP1 and CYFIP2 are 87% identical at the amino acid level and no difference concerning Rac binding or function within WRC could hitherto be established for the two isogenes. An alignment of multiple CYFIP variants found in various experimental model systems down in evolution from chicken, fish and fly till slime mold (*D. discoideum*) and plants revealed, firstly, that all residues mutated in CYFIP2 of human patients are conserved in CYFIP1 of human, mouse, chicken and fish, and secondly, that in spite of very few exceptions, the degree of conservation in mutant residues was striking, down to species as distant as *Dictyostelium discoideum* (Figure S1). Notably, our conclusions on the differential regulation of murine CYFIP1 by its two distinct Rac binding sites was fully recapitulated by the same mechanisms in the single *pirA* gene in *Dictyostelium discoideum*, strongly suggesting that respective mechanisms are fundamental and relevant across species barriers [7].

**Figure 1.**
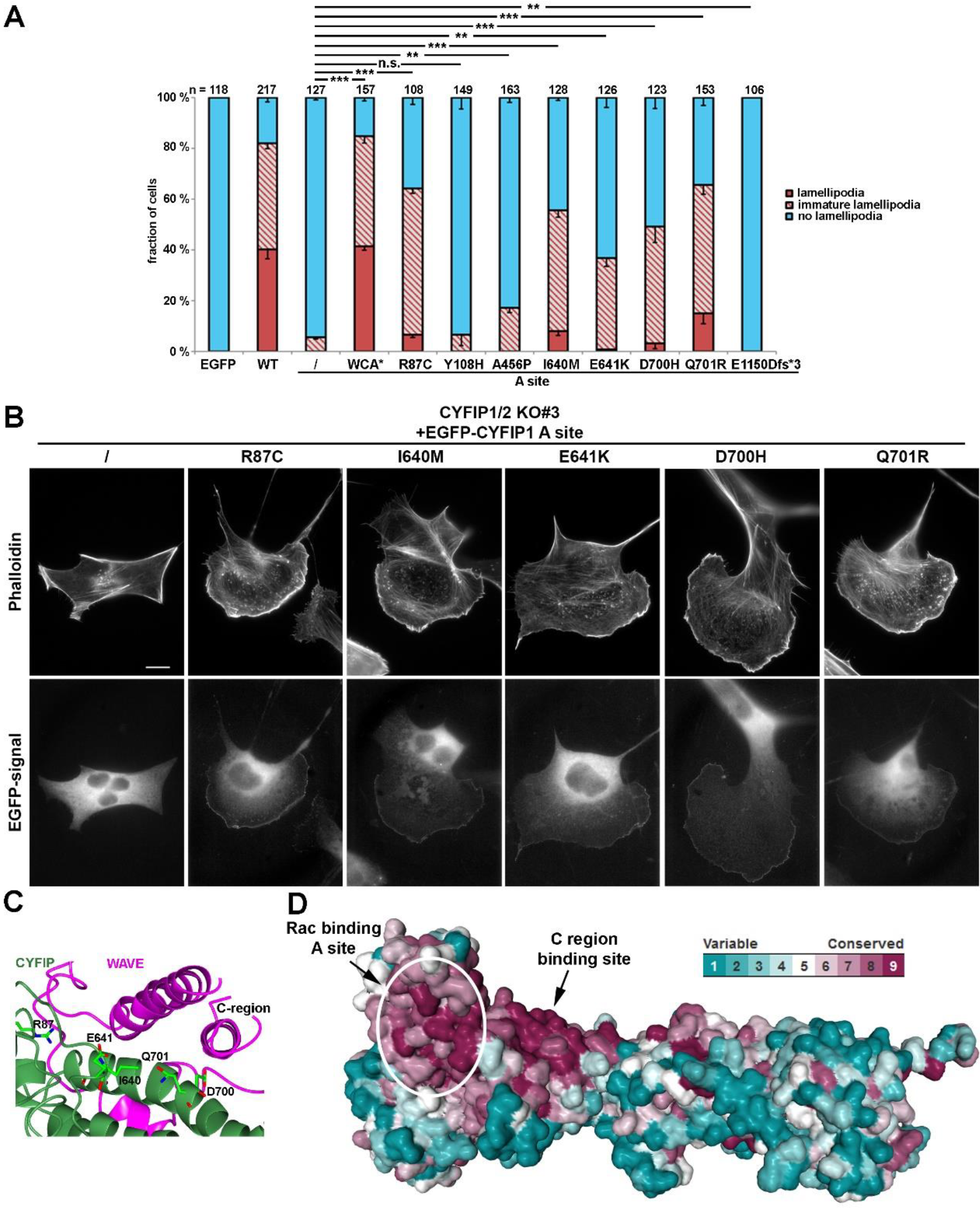
Intellectual disability-causing mutations in CYFIP frequently cause constitutive activation of WRC. (A) Quantification of lamellipodia formation. Indicated EGFP-tagged CYFIP1 constructs (or EGFP as control) were transfected into B16-F1 CYFIP1/2 KO cells (clone #3) and lamellipodia formation was assessed after staining of the actin cytoskeleton with ATTO 594-labeled phalloidin. Lamellipodial actin networks that were generally small, narrow, irregular or displayed multiple ruffles were defined as ‘‘immature lamellipodia’’, as opposed to regular, fully developed lamellipodia. n gives number of cells analyszed, data correspond to arithmetic means ± SEM from at least three independent experiments. Statistical significance was assessed for differences between the percentages of cells with “no lamellipodia” phenotype. n.s.: not statistically significant; **p<0.01; ***p<0.001 (two-sample, two-sided t-test). (B) Cell morphologies of CYFIP1/2 KO cells (clone #3) expressing respective constructs, as indicated. Panels in top row show stainings of the actin cytoskeleton with phalloidin, and bottom row images show fluorescence of the same cells derived from EGFP fluorescence. (C) Close up view of the interface between CYFIP and the C-region of WAVE, required for transinhibition of WAVE’s WCA domain. Mutations in CYFIP presumably causing constitutive activation of WRC are shown. (D) Surface conservation of CYFIP. Note that high sequence conservation is found in a larger area contacting the C-region of WAVE in which activating mutations are located.

Introduction of ID-causing CYFIP2 mutations into the A site-mutated CYFIP1 background revealed that constitutive activation of WRC can also be achieved by mutations alternative to WCA*. While the A site mutant was capable of triggering the formation of at best immature lamellipodia, and only at very poor frequency (<5% of transfected cells, Figure 1A), additional mutation of CYFIP2 residues mutated in patients suffering from ID, specifically R87C, I640M, E641K, D700H and Q701R (positions in CYFIP1) restored lamellipodia formation to substantial extent, albeit not quite as completely as seen with A site-mutated WCA* (Figures 1A, B). Restoration of lamellipodia formation by these constructs also coincided with their robust accumulation at the tips of lamellipodia now formed, a subcellular localization not seen for the mostly cytosolic, EGFP-tagged A site mutant of CYFIP1 (Figure 1B, left panels). However, since the rescue of lamellipodia formation was not as strong as disrupting the WH2 (W)- and C-region contact sites in CYFIP in the WCA* variant, our data indicated a partial rather than full activation of WRC by these patient-derived mutations.

Interestingly, all these residues mutated in patients either directly contact the C-region of WAVE or are positioned at least in close proximity to the C region binding site (Fig 1C). Investigating the evolutionary surface conservation indicates that the C-region binding site of CYFIP is extremely highly conserved (Figure 1d), potentially explaining why modifications in this region result in impaired C-region binding, which is necessary for WCA domain sequestration [10]. Interestingly, the A456P mutation only caused a slight, yet statistically significant increase in lamellipodia formation in our conditions. Moreover, the Y108H mutation did not show any apparent effect in our assay, although it was previously also proposed in regulating WRC activation [24].

In our attempts to further characterize the activating potential of specific ID-causing CYFIP mutations, we initially focused on R87C, as this mutant was reported to occur recurrently, in two independent studies [23,24], and caused a robust induction of lamellipodia formation in the context of abrogated A site function (Fig 1A, B). Introduction of the R87C mutation or WCA* into WT CYFIP1 did not change the frequency of lamellipodia formation upon rescue in CYFIP1/2 KO cells (clone #3) (Figure S2). This was simply due to the fact that Rac-mediated WRC activation in B16-F1 cells migrating on laminin is already at an optimum, and not apparently the limiting factor if restoration of WRC function is undertaken with wildtype CYFIP1 [7]. A potential improvement of activation as seen in the context of the A site mutation can thus not be phenotypically documented in wildtype CYFIP1 conditions.

However, to illustrate further that the R87C mutation enhances the activation state of WRC *in cellulo*, we also sought to employ a complementary approach, which was independent of introduction of additional mutations in respective CYFIP subunit, as occurring in patients. To do this, we employed a B16-F1 melanoma clone that had arisen from our attempts to disrupt Rac1/2/3 genes in the CYFIP1/2 null background, as described previously [26]. The novel clone (termed CYFIP1/2 KO#3+Rac1/2/3 KO#4; see Figure 2A), hitherto unpublished, which was also selected upon CRIPSR/Cas9-treatment targeting all Rac alleles, displayed reduced, but not entirely abolished Rac expression, as seen in previously published #11 (Figure 2A, [26]). We hypothesized that if reduced Rac expression levels were indeed physiologically relevant for the induction of lamellipodia formation in this cell line, rescue with WT CYFIP1 would cause a reduced frequency of this response as compared to the parental CYFIP1/2 null clone (see Figure 1), and this was indeed the case. Lamellipodia formation upon rescue with WT CYFIP1 was observed to be below 20%, but could be significantly improved in frequency by additional R87C or again, WCA* mutations (Figure 2B). Moreover, in those WRC-deficient, low Rac expressing cells (CYFIP1/2 KO#3+Rac1/2/3 KO#4), in which lamellipodia were formed upon rescue with R87C or WCA*, these lamellipodia often accumulated WRC at their tips at increased intensities than observed for WT CYFIP1 (Figure 2C). To explore more directly if increased lamellipodia formation frequency (Figure 2B) and WRC accumulation intensity (Figure 2C) was also able to translate into increased actin assembly efficiency, we examined the velocity of protrusion of the cell edges formed in different conditions. We had previously established a clear correlation between WRC gene dose and rate of protrusion or lamellipodial Arp2/3 complex activation ([27]; for review see also [2]), so changes in capability to activate WRC should also be reflected in protrusion efficacy. And indeed, whereas WT CYFIP1 on average failed to promote cell edge protrusion beyond the levels observed in B16-F1 lacking endogenous WRC and reduced for Rac expression, R87C as well as WCA* mutants in these cells led to an increase in protrusion speed by at least two fold. This strongly suggests that a subset of ID-causing CYFIP2 mutations give rise to WRCs that can more easily be activated or are already partially activated.

**Figure 2.**
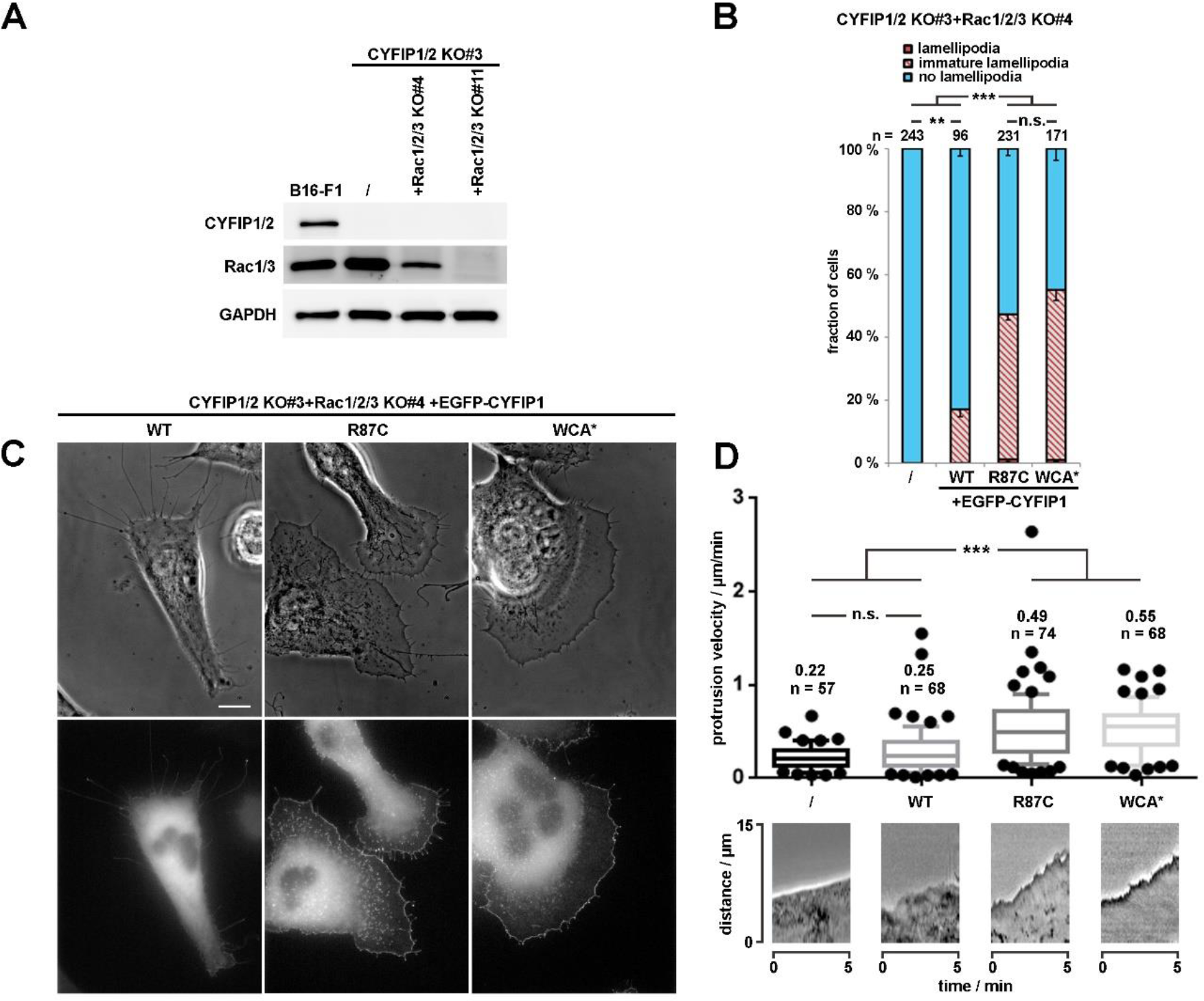
R87C mutation in CYFIP enhances lamellipodium protrusion at low Rac GTPase levels. (A) Western blotting of distinct cell lines to probe for expression levels of endogenous CYFIP and/or Rac GTPases. (B) Quantification of lamellipodia formation in B16-F1 CYFIP1/2+Rac1/2/3 KO cells (clone #3/4) harbouring low Rac expression and transfected with indicated EGFP-tagged CYFIP1 constructs, as described in Figure 1A. (C) Live cell imaging of CYFIP1/2+Rac1/2/3 KO cells (clone #3/4) expressing EGFP-tagged CYFIP1 constructs. Upper panels show phase contrast images, and bottom panels EGFP fluorescence. (D) Quantification of cell edge advancement. n gives the number of cells analysed. Box and whisker plots represent data as follows: boxes correspond to 50% of all data points (25%–75%), and whiskers to 80% (10%–90%). Lines and numbers above boxes correspond to medians. Statistical significance was assessed by non-parametric, Mann-Whitney-Rank-Sum test, n.s.: not statistically significant; ***p<0.001. Bottom part shows representative kymographs.

As opposed to R87C and most other patient-derived mutations, the truncating mutation E1150Dfs*3 caused a complete failure in triggering lamellipodia formation in the A site-mutated CYFIP1 (Fig 1A), which already indicated that this mutation cannot likely cause a variant that is constitutively activated. To directly test its capability of rescuing lamellipodia formation in CYFIP1/2 null cells, the modifications were introduced into the background of WT CYFIP1, and explored upon transfection into B16-F1 lacking endogenous WRC (CYFIP1/2 KO#3, [7]). Single cell analyses revealed the absence of rescue of lamellipodia formation with this variant (Figure 3A, B). Immunoblotting using bulk cell extracts of CYFIP1/2 null cells transiently transfected with WT CYFIP1, R87C- or E1150Dfs*3-CYFIP1 indicated that the latter variant is only weakly expressed as compared to WT CYFIP1 or R87C (Figure 3C). Moreover, E1150Dfs*3 mutant was unable to assemble into endogenous WRC, as it could not co-precipitate endogenous WRC subunits Nap1, WAVE and Abi1 in wildtype B16-F1 cells, although all of them were readily immunoprecipitated by EGFP-tagged WT or R87C-CYFIP1 (Figure 3D). The inability of the E1150Dfs*3 variant to associate with remaining WRC subunits then also explains its incapability to rescue the well-established downregulation of WAVE expression in CYFIP1/2 KOs, again as opposed to WT CYFIP1. Together, these data reveal that E1150Dfs*3 constitutes a “null” allele unable to associate with remaining WRC subunits and thus failing to operate as functional unit linking WRC to Rac signalling.

**Figure 3.**
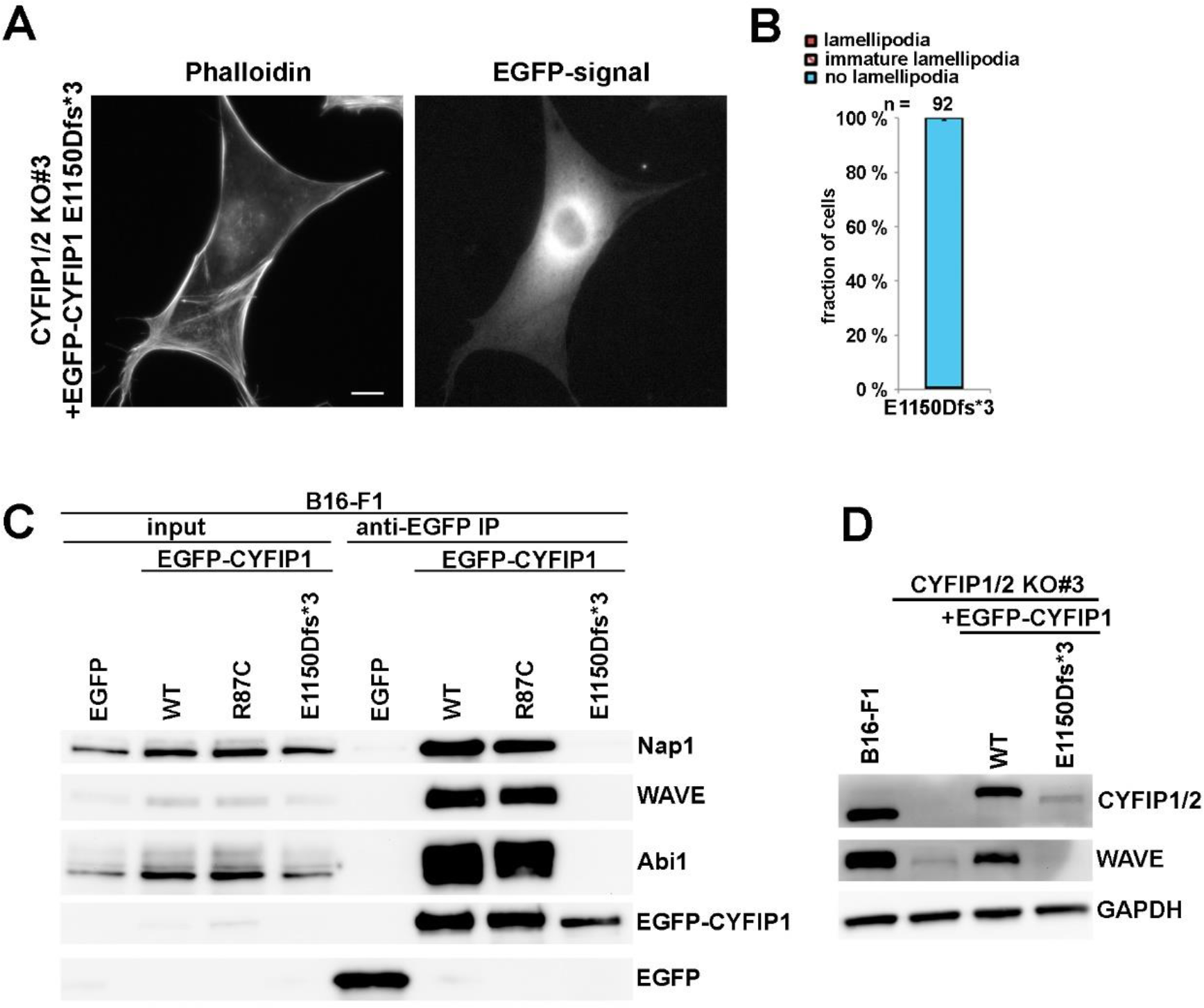
C-terminal truncation of CYFIP prevents WRC assembly and lamellipodia formation. (A) Cell morphology of CYFIP1/2 KO cells (clone #3) expressing EGFP-tagged E1150Dfs*3 CYFIP1 mutant, leading to a C-terminal truncated protein. (B) Quantification of lamellipodia formation, as described in Figure 1A. Note that CYFIP1 WT control is as in Fig. 1A. (C) B16-F1 cells were transfected with indicated constructs, lysed and subjected to immunoprecipitation analysis to assay interaction of E1150Dfs*3 mutant CYFIP1 with WRC components WAVE and Nap1. (D) Western blotting of B16-F1 cells or CYFIP1/2 KO cells (clone #3) transfected with indicated constructs and probed for expression of CYFIP and WAVE.

## 4. Discussion

In this study, we systematically analysed CYFIP2 mutations, reported to cause intellectual disability [23,24]. WRC is a highly conserved protein complex. We could previously show that the regulation of WRC by Rac GTPase is conserved from *D. discoideum* to mammals [7]. Furthermore, both Rac binding sites, as well as the WH2- and C-region contact sites of WAVE are conserved in CYFIP1 and CYFIP2, strongly suggesting that their basic regulation by Rac are indistinguishable. In this study, we show that CYFIP mutations positioned within or in close proximity to the C region binding site behave similar to constitutively active WRC. Upon abrogation of the Rac binding A site, which is crucial for allosteric activation of WRC, CYFIP1 mutations R87C, I640M, E641K, D700H, Q701R can all rescue lamellipodia formation, in a fashion comparable to the known constitutively active mutant WCA* [7,10]. In a complementary assay, in which cellular Rac levels were reduced in the absence of endogenous CYFIP, the R87C mutation behaved comparable again to WCA* concerning its capability of rescuing lamellipodia formation.

Although aforementioned mutations likely promote WRC activation, a mutation causing truncation of the C-terminal 100 amino acids (E1150Dfs*3) generates a loss-of-function variant, entirely abolished for assembling into WRC and thus driving lamellipodia formation. This appeared puzzling at first glance. However, WAVE1 mutations causing partial WCA domain truncations and thus behaving as dominant negatives have also been shown to be causative for ID [21]. This thus suggests that a precisely tuned, narrow window of WRC activity in the human brain is required for its proper development and function, with both hyper-activation and downregulation leading to similarly drastic consequences. Supporting this hypothesis, a recent study also suggested that tight regulation of Rac activity is required for proper neurological function, and that both hyper- and hypoactivation of Rac signalling can cause ID [28]. However, both C-terminally truncated and thus dominant-negative WAVE1 mutants and activating CYFIP2 mutants will likely display enhanced Rac binding when assembled into WRC, at least in theory. More specifically, it is well established that experimental interference with the CYFIP-WCA domain interaction dramatically enhances Rac binding, since Rac binding-mediated WRC activation competes with the CYFIP-WCA domain interaction and mutating the WCA domain contact sites in CYFIP and deleting WAVE’s WCA domain causes the same effect with respect to Rac binding [10]. It is possible therefore that both WAVE truncation and activating CYFIP2 mutations can similarly sequester away active Rac, thereby suppressing Rac functions and its effector pathways. This could work in a fashion similar to what has recently been suggested for the Rac binding protein FAM49B [29–31]. Such a scenario would also fit the observation that certain missense mutations in Rac1 leading to ID cause its loss of function [22], as well as the model that RhoA signalling is typically upregulated and Rac signalling suppressed in ID [32].

Interestingly, we failed to observe an activating effect by the Y108H mutation in spite of previous proposals [24]. This suggests that Y108H is likely not causing major alterations of WRC structure. However, publicly available databases indicate Y108 to constitute the by far most phosphorylated residue in CYFIP proteins (www.phosphosite.org). It is possible therefore that Y108 phosphorylation of CYFIP2 is crucial for specific, neurodevelopmental processes that cannot be covered by our *in cellulo* assay designed to examine sole changes in WRC activity. Although this remains to be confirmed in future studies, the data already suggest that even in neurons, the changes caused by the Y108H-mutation will not likely involve modulation of WRC activity.

It was previously suggested that WRC activation occurs via disassembly of the complex, leading to a CYFIP/Nap/Abi and a WAVE/HSPC300 subcomplex, respectively [11] and that ID-related CYFIP2 mutations would act by destabilising WRC, favouring disassembly [24]. This view, however, is challenged by various observations indicating that upon WRC activation, the complex stays intact [7,9,10]. In line with the latter view, we found the activating mutation R87C to normally interact with WRC components, while the truncating mutation E1150Dfs*3 that fails to cause lamellipodia formation to show no interaction with WRC components. All these data are in accordance with a revised model of WRC activation, in which activation occurs through release of WCA-domain for Arp2/3 complex activation and actin filament branching, while the WRC unit remains intact [7,9,10]. Taken together, our results demonstrate that ID-causing CYFIP2 mutations can have opposing effects on WRC activity, in contrast to what has been anticipated previously [24], and emphasize the need for robust and clean experimental readouts, such as CRISPR/Cas9-based gene disruption and rescue.

## Supplementary Materials

Figure S1: Sequence alignment of CYFIP paralogs, Figure S2: Lamellipodia formation by constitutively active WRC.

## Author Contributions

Conceptualization and methodology, M.S.; investigation, M.S.; resources, K.R.; writing—original draft preparation, M.S.; writing—review and editing, K.R.; supervision, K.R.; funding acquisition, K.R. All authors have read and agreed to the published version of the manuscript.

## Acknowledgments

We would like to thank all members of the lab for continuous and fruitful discussions.

