## Supplementary figures for "Molecular dissection of neurodevelopmental disorder-causing mutations in CYFIP2"

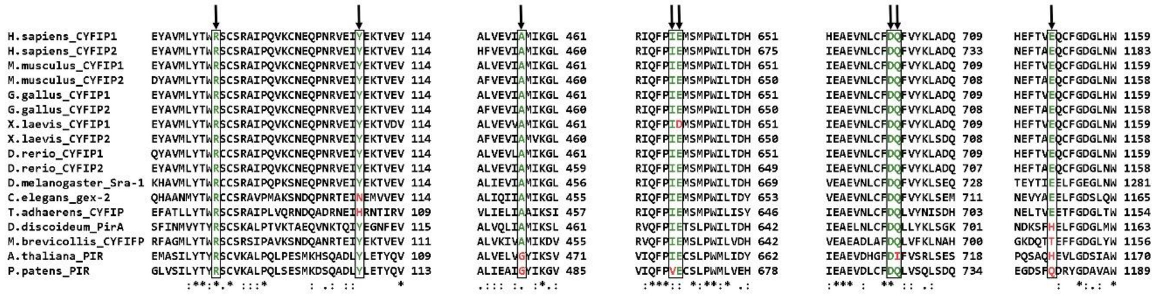


**Figure S1: Sequence alignment of CYFIP paralogs.** A multiple sequence alignment was performed on CYFIP paralogs. Intellectual disability-related CYFIP2 mutations are highlighted. Amino acid conservation or divergence during evolution is indicated by green and red colours, respectively.

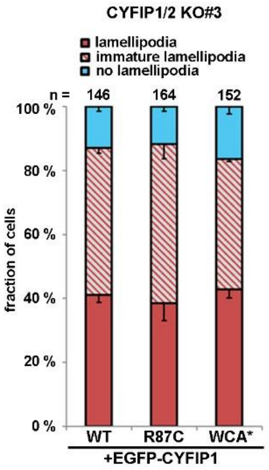


**Figure S2: Lamellipodia formation by constitutively active WRC.** ) Quantification of lamellipodia formation in B16-F1 CYFIP1/2 (clone #3) transfected with indicated EGFP-tagged CYFIP1 constructs, as described in Figure 1A.
